# Multiscale model of heart growth during pregnancy: Integrating mechanical and hormonal signaling

**DOI:** 10.1101/2020.09.18.302067

**Authors:** Kyoko Yoshida, Jeffrey J. Saucerman, Jeffrey W. Holmes

## Abstract

Pregnancy stands at the interface of mechanics and biology. The growing fetus continuously loads the maternal organs as circulating hormone levels surge, leading to significant changes in mechanical and hormonal cues during pregnancy. In response, maternal soft tissues undergo remarkable growth and remodeling to support the mother and baby for a healthy pregnancy. We focus on the maternal left ventricle, which increases its cardiac output and mass during pregnancy. This study develops a multiscale cardiac growth model for pregnancy to understand how mechanical and hormonal cues interact to drive this growth process. We coupled a cell signaling network model that predicts cell-level hypertrophy in response to hormones and stretch to a compartmental model of the rat heart and circulation that predicts organ-level growth in response to hemodynamic changes. We calibrated this multiscale model to data from experimental volume overload (VO) and hormonal infusions of angiotensin 2 (AngII), estrogen (E2), and progesterone (P4). We then validated the model’s ability to capture interactions between inputs by comparing model predictions against published observations for the combinations of VO+E2 and AngII+E2. Finally, we simulated pregnancy-induced changes in hormones and hemodynamics to predict heart growth during pregnancy. Our model produced growth consistent with experimental data. Overall, our analysis suggests that the rise in P4 during the first half of gestation is an important contributor to heart growth during pregnancy. We conclude with suggestions for future experimental studies that will provide a better understanding of how hormonal and mechanical cues interact to drive pregnancy-induced heart growth.

## 1. Introduction

Pregnancy stands at the interface of mechanics and biology. Throughout nine months of pregnancy, the growing fetus continuously loads the maternal organs while circulating hormone levels surge. Pregnancy presents a cardiovascular challenge to the mother as her cardiac output increases by 45%, her total systemic resistance falls by 34%, and her total blood volume increases by almost 50% during a normal, singleton pregnancy (Pritchard 1965; Hunter and Robson 1992). Simultaneously, circulating estrogen (E2) and progesterone (P4) hormone concentrations increase up to 10 times above nonpregnant levels (Tulchinsky et al. 1972). In response to these dynamic mechanical and hormonal cues, maternal soft tissues undergo significant growth and remodeling to support both mother and baby. We focus here on the maternal left ventricle (LV), which increases in mass and cavity volume by approximately 30% during the course of a normal pregnancy and regresses to normal size between 3-6 months after delivery (Savu et al. 2012).

Cardiovascular conditions and cardiomyopathy together account for more than 25% of pregnancy-related deaths (Creanga et al. 2017). With maternal death rates in the United States rising over the last 25 years (Kassebaum et al. 2016), there is a critical need to understand how pregnancy hormones and hemodynamic changes interact to affect heart growth. Cardiac hypertrophy is a complex, multiscale process, driven by both mechanical and hormonal stimuli that interact through an extensive network of intracellular pathways within a cardiac muscle cell (cardiomyocyte). Experimentally, hypertrophy can be triggered by either hemodynamic or pharmacologic interventions. An increase in aortic resistance due to constriction (pressure overload, PO) leads to mass increase and wall thickening, while an increase in cardiac output due to an arteriovenous fistula or valvular regurgitation (volume overload, VO) leads to mass increase and LV cavity dilation. Infusion of isoproterenol leads to LV mass increase (De Windt et al. 2001; Drews et al. 2010), as does infusion of angiotensin (Griffin et al. 1991; Dostal and Baker 1992). Pregnancy hormones also influence heart growth. *In vivo* and *in vitro* studies have shown the ability of E2 to attenuate growth (Pedram et al. 2008), and the role of P4 in promoting growth (Chung et al. 2013). Furthermore, mechanical and hormonal stimuli can interact to impact growth, as demonstrated by attenuated LV mass increase in experiments of volume and pressure overload in E2 treated animals compared to untreated controls (Voloshenyuk and Gardner 2010; Iorga et al. 2016).

Computational models have made considerable progress in predicting heart growth in response to both hormonal and mechanical stimuli (Yoshida and Holmes 2020). Computational systems biology models can predict cell-level cardiomyocyte growth in response to hormones and pharmaceutical interventions (Ryall et al. 2012; Frank et al. 2018). Mechanics-based growth models can predict heart growth at the organ level in multiple settings where changes in mechanical loading of the heart occur, including PO, VO, myocardial infarction, and dyssynchrony (Witzenburg and Holmes 2017, 2018). Recently, our group developed a multiscale model that incorporates mechanical and hormonal stimuli to predict concentric heart growth in response to pressure overload (Estrada et al. 2020). The objective of this work is to build a multiscale cardiac growth model for pregnancy by combining a systems biology model with a mechanics-based model to capture the interactions between hormonal and mechanical stimuli. Towards this objective, we created a cell signaling network model that predicts cell-level growth in response to the pregnancy hormones E2, P4, as well as angiotensin 2 (AngII), and stretch. We coupled this network model to a compartmental model of the rat heart and circulation to capture changes in hemodynamics. Because pregnancy is associated with both VO and elevated hormones, we first calibrated this multiscale model to quantitatively match experimental heart growth in response to VO and infusions of AngII, E2, and P4. We then validated model predictions against experimental observations of heart growth in response to the combinations of AngII+E2 and VO+E2. Finally, we simulated pregnancy changes in hormones and hemodynamics to predict heart growth during pregnancy. Our multiscale model was able to capture reported pregnancy changes in both LV growth and hemodynamics. Furthermore, our analysis suggests that heart growth during pregnancy is in part driven by the early rise in P4, particularly during the first half of gestation.

## 2. Materials and Methods

### 2.1 Intracellular signaling network model of cardiomyocyte growth

To simulate the influences of hormones and stretch on cell-level growth, we built an intracellular signaling network model for cardiomyocyte hypertrophy (Fig. 1A). Our approach was similar to a model developed by Ryall et al. (Ryall et al. 2012; Frank et al. 2018) that included 107 species interacting through 207 reactions representing the various intracellular hypertrophic pathways. Since the Ryall model did not include pregnancy hormones, and since limited experimental data on the effect of P4 on cardiomyocyte hypertrophic pathways are available (Chung et al. 2012, 2013), we focused on a subset of species within the original network model for which experimental data are available. Our network (Fig. 1A) consisted of 4 key inputs: the pregnancy hormones Progesterone (P4) and Estrogen (E2); Angiotensin 2 (AngII), another hormone elevated in response to both volume overload (Ruzicka and Leenen 1995) and pregnancy (Mishra et al. 2018); and Stretch, a generic stimulus reflecting mechanotransduction pathways that activate Akt and MEK/ERK signaling. These inputs interacted with each other and nine other intermediate species through 24 reactions, which led to changes in cardiomyocyte size, represented by the output, CellArea.

**Figure 1:**
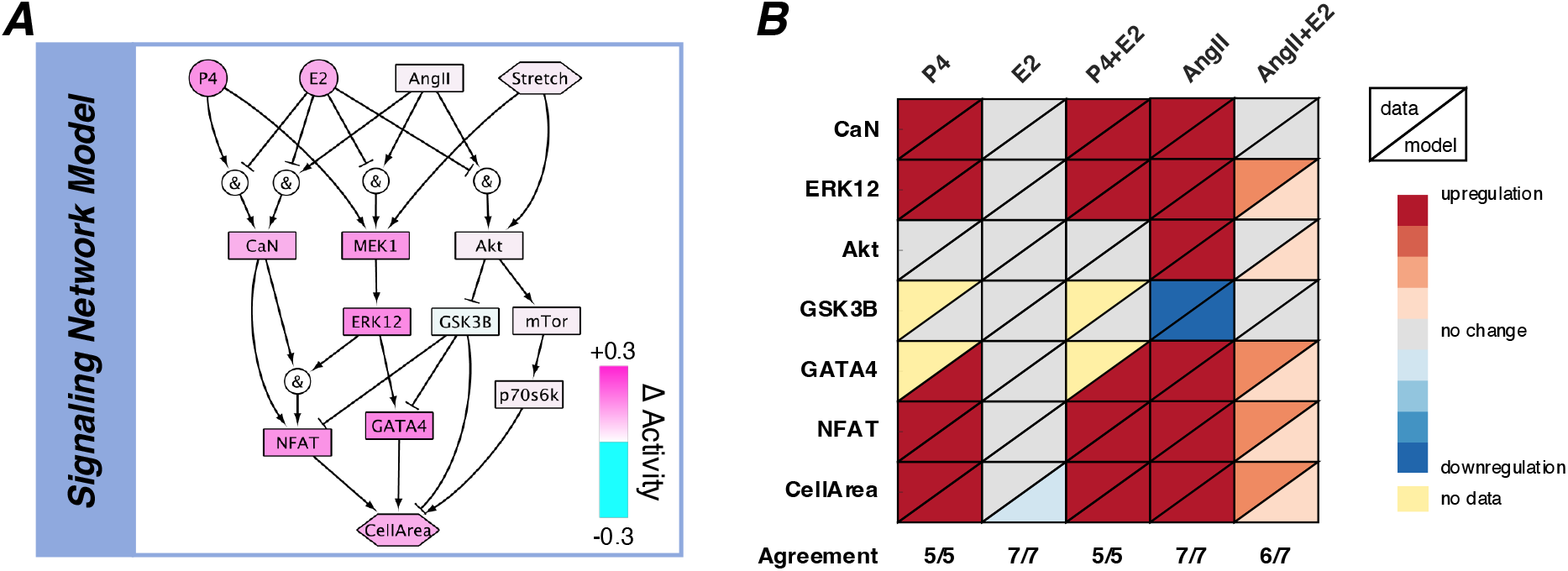
Intracellular signaling network model. A.) The intracellular signaling network model of cardiomyocyte growth predicts the amount of growth due to a particular level of hormones (P4, E2, AngII) and Stretch, and outputs a CellArea. Colors represent example activity levels at the end of pregnancy, where pink and blue represent increased and decreased activity from baseline respectively. B.) Network model calibration against *in vitro* data. Upper left triangles represent phosphorylation or activity reported from experiments where neonatal cardiomyocytes were treated with P4, E2, P4+E2, AngII, or AngII+E2 (columns). Red represents upregulation, grey represents no change, and blue represents downregulation of activity for each signaling intermediate (rows). Yellow color signifies no available data for a particular species and experimental condition. Lower right triangles represent network model predictions for each experimental condition (More details and experimental study references are available in Supplemental Data 2: InVitroCalibration).

We modeled reactions in the network using normalized Hill-type ordinary differential equations in which the level of activation of each species is represented as a normalized value between 0 and 1, where 0 indicates no signaling activity, and 1 represents maximal activation (Kraeutler et al. 2010). Equation (1) shows an example of the differential equation for the intermediate species, Akt, which is activated by AngII or Stretch, but inhibited by E2 in combination with AngII activation:

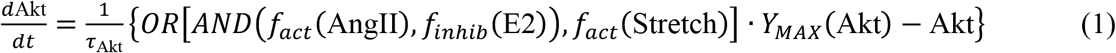

Here, τ_Akt_ is the species time constant and *Y*_*MAX*_ is the maximal fractional activation of Akt. OR and AND represent logical operations, which describe crosstalk between species:

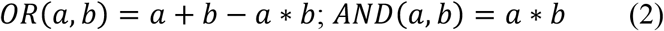

The activation/inhibition functions in Equation (1) are defined such that *f*_*act*_(0) = 0, *f*_*act*_(1) = 1, *f*_*inhib*_(0) = 1, and *f*_*inhib*_(1) = 0:

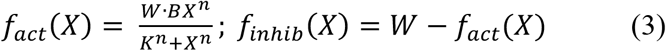

where *W* is the reaction weight, *n* is the Hill coefficient, and the functions *B* and *K* are defined as:

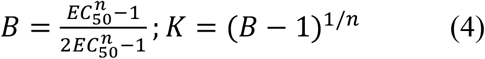

where *EC*_*50*_ is the half-maximal activation. The complete list of equations is available in the Supplemental Document. The MATLAB code for this model (E2P4HypertrophyNetwork.m) is available on SimTK (https://simtk.org/projects/heartgrowthpreg) and was automatically generated from Supplemental Data 1 using the freely available software Netflux (https://github.com/saucermanlab/Netflux).

We simulated changing levels of hormones and stretch by adjusting the reaction weights of the inputs (*w*_AngII_, *w*_E2_, *w*_P4_, *w*_Stretch_) based on circulating hormone concentrations and calculated stretch as described below (Section 2.3). We set the time constant, τ, to 0.1 hours for all species except CellArea, which we set to τ_CellArea_ = 300 hours to represent the longer time necessary for protein accumulation and cell growth compared to phosphorylation events. For all other parameters, (*Y*_max_, *EC*_50_, *n*) we used the same default values as in Ryall et al. (Ryall et al. 2012) (Table 1). We based reactions for Stretch and AngII on Ryall et al., and adjusted crosstalk interactions (AND/OR) for E2 and P4 to match phosphorylation data from *in vitro* experiments where neonatal rat ventricular cardiomyocytes were treated with P4 alone, E2 alone, AngII alone, or the combinations of P4+E2 and AngII+E2 (Fig. 1B, Supplemental Data 2: InVitroCalibration). The final version of the network model correctly matched reported trends for 30 out of 31 phosphorylation data from 5 different publications (Pedram et al. 2005, 2013a, b; Chung et al. 2012, 2013). The one disagreement between the model and experimental data was for Akt activity under the combination of AngII+E2. The model predicted a slight increase in Akt activity, while no significant difference was reported in the experimental study (Pedram et al. 2005).

**Table 1:** Signaling network model parameters

| Parameter | Baseline value |
| --- | --- |
| Reaction weight ($W$) | 0.1335 for inputs ($W_{Stretch}$, $W_{AngII}$, $W_{E2}$, $W_{P4}$)<br>1 for all other species |
| Maximal activation ($Y_{max}$) | 1 |
| Half-maximal activation ($EC_{50}$) | 0.5 |
| Hill coefficient ($n$) | 1.4 |
| Reaction time constant ($\tau$) | 0.1 hr Input and Middle Species<br>300 hr CellArea Output |

### 2.2 Compartmental model of organ-level growth

Our group previously published a rapid, compartmental growth model of the heart coupled to a lumped-parameter circuit model based on a canine anatomy, and demonstrated its ability to predict heart growth in multiple settings where changes in mechanical loading of the heart occur, including volume overload (VO), pressure overload (PO), and myocardial infarction (Santamore and Burkhoff 1991; Witzenburg and Holmes 2018). Briefly, this model uses a time-varying elastance model to simulate the ventricles. The end-systolic pressure-volume relationship is defined by a linear elastance and the end-diastolic pressure-volume relationship is defined by an exponential function. The systemic and pulmonary aorta and vessels are represented by capacitors in parallel with resistors (Fig. 2A, Supplemental Table 1), and the stressed blood volume (*SBV*) is defined as the total blood volume contained in these capacitors and ventricles. This model is implemented as a system of differential equations in MATLAB to simulate changes in the volume of each compartment over a cardiac cycle. This code is freely available to download on SimTK (https://simtk.org/projects/heartgrowthpreg).

**Figure 2:**
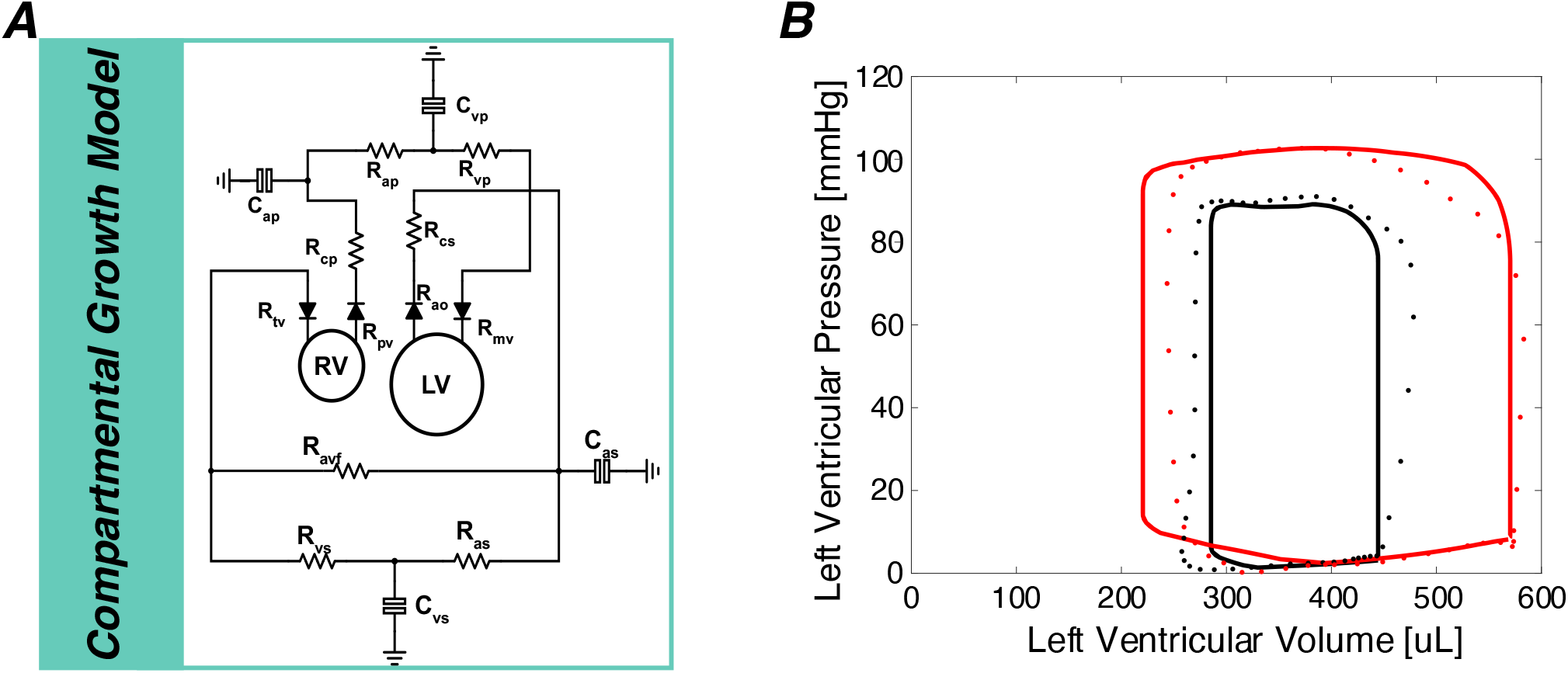
A.) Compartmental model of organ-level growth. calculates changes in elastic stretches (*F*_*e*_) in the left ventricle due to growth (*F*_*g*_) and hemodynamic alterations. The right and left ventricles are modeled as thin-walled spheres. The resistors and capacitors represent the resistance and compliance of the systemic and pulmonary arteries and veins while the diodes represent the valves and allows flow only in one direction (Table 2 and Supplemental Table 1). B.) Pressure volume behavior for baseline and acute volume overload generated by the calibrated compartmental model. Dots represent pressure-volume data points from (Holmes 2004) and solid lines represent model fit to the data. Black: Baseline, Red: acute volume overload.

#### 2.2.1 Simulating baseline and acute volume overload hemodynamics

Because experimental studies of VO, hormonal infusion, and pregnancy are more common in small animals, we reparametrized the canine compartmental model to match control hemodynamics from rats using allometric scaling methods (West et al. 1997; Holmes 2004; Caggiano et al. 2022) (See Supplemental Document for more details). To simulate volume overload in rats induced via an arteriovenous fistula between the abdominal aorta and inferior vena cava (Garcia and Diebold 1990), we added a resistor to represent the fistula (*R*_*avf*_) between the aorta and the vena cava (Fig. 2A). To calculate stretches experienced by the heart during baseline and acute VO states, we optimized the circulatory parameters to match averaged experimental hemodynamics reported by Holmes et al. for control rats and for rats subjected to acute VO (4 days post-fistula) (Holmes 2004).

We specified the reported heart rate (*HR*) and the maximum slope of the end-systolic pressure-volume relationship (*E*) directly in the model for both baseline and acute VO. We adjusted end-diastolic pressure-volume parameters (*A, B, V*_*0*_) to match experimental pressures and volumes during the filling phase of the cardiac cycle. Baseline systemic resistance (*R*_*as*_) and stressed blood volume (*SBV*) were adjusted to match the reported control values of end-systolic pressure (ESP) and end-diastolic volume (EDV). Holding the baseline *R*_*as*_ constant, the fistula resistance (*R*_*avf*_) and *SBV* were adjusted to match experimental ESP and EDV 4 days after fistula creation. All parameters were simultaneously optimized using the fminsearch function in MATLAB, until the differences between measured and predicted hemodynamic values were minimized (Witzenburg and Holmes 2018). To confirm the uniqueness of the optimized circulation model parameters, we conducted a Monte Carlo simulation (Supplemental Document, Suppl. Fig. 1). Figure 2B illustrates the agreement between the final calibrated baseline and acute VO model to the experimental pressure-volume behavior, and Table 2 summarizes the resulting circulation model parameters and hemodynamic comparisons between the model and data.

**Table 2:**
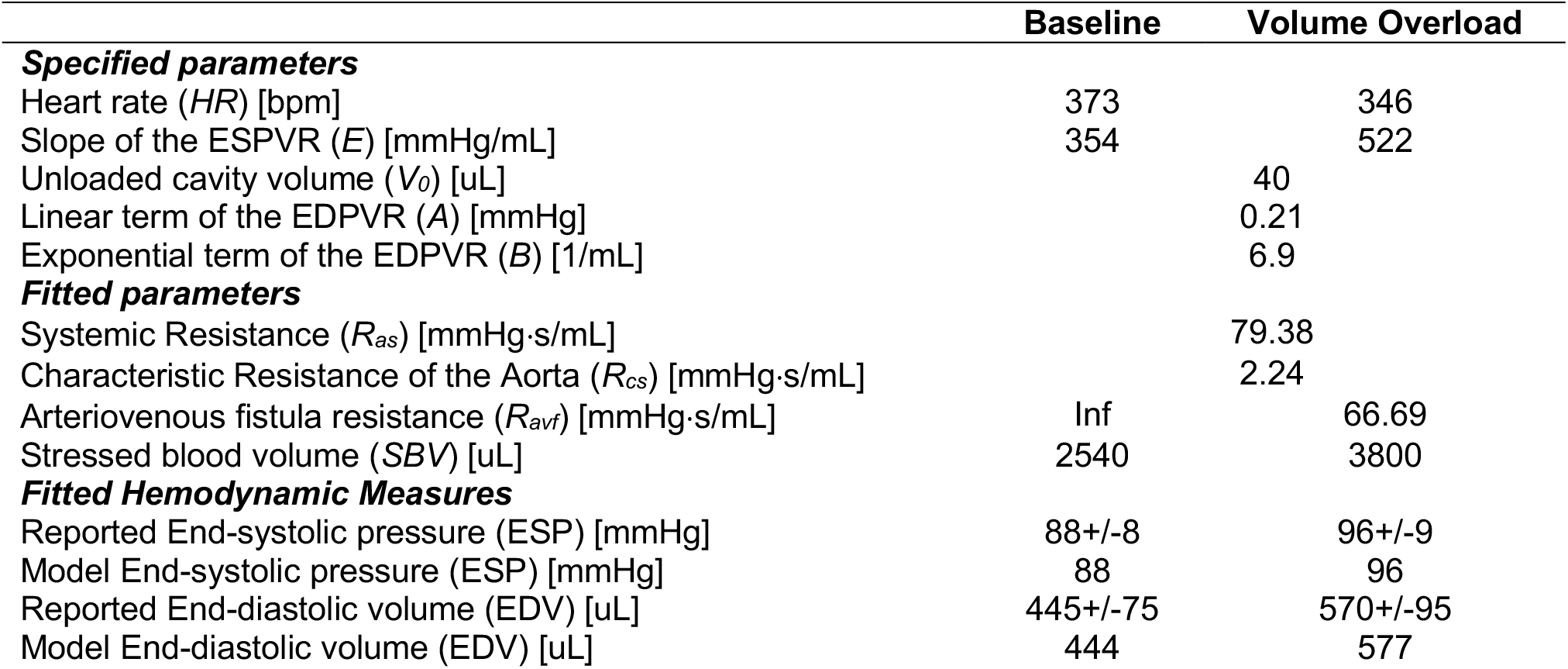
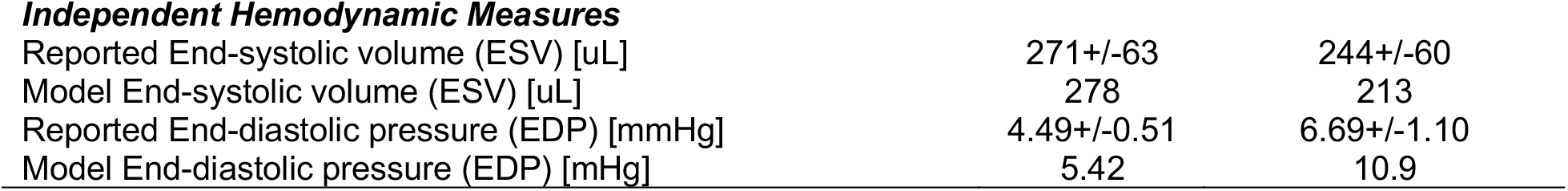
Model parameters for volume overload simulations. Reported hemodynamic data are from (Holmes 2004). ESPVR: end-systolic pressure-volume relationship and EDPVR: end-diastolic pressure-volume relationship.

|  | Baseline | Volume Overload |
| --- | --- | --- |
| **Specified parameters** |  |  |
| Heart rate ($HR$) [bpm] | 373 | 346 |
| Slope of the ESPVR ($E$) [mmHg/mL] | 354 | 522 |
| Unloaded cavity volume ($V_0$) [μL] | | 40 |
| Linear term of the EDPVR ($A$) [mmHg] | | 0.21 |
| Exponential term of the EDPVR ($B$) [1/mL] | | 6.9 |
| **Fitted parameters** |  |  |
| Systemic Resistance ($R_{as}$) [mmHg·s/mL] | | 79.38 |
| Characteristic Resistance of the Aorta ($R_{cs}$) [mmHg·s/mL] | | 2.24 |
| Arteriovenous fistula resistance ($R_{avf}$) [mmHg·s/mL] | Inf | 66.69 |
| Stressed blood volume ($SBV$) [μL] | 2540 | 3800 |
| **Fitted Hemodynamic Measures** |  |  |
| Reported End-systolic pressure (ESP) [mmHg] | 88+/-8 | 96+/-9 |
| Model End-systolic pressure (ESP) [mmHg] | 88 | 96 |
| Reported End-diastolic volume (EDV) [μL] | 445+/-75 | 570+/-95 |
| Model End-diastolic volume (EDV) [μL] | 444 | 577 |
| Independent Hemodynamic Measures |  |  |
| Reported End-systolic volume (ESV) [uL] | 271+/-63 | 244+/-60 |
| Model End-systolic volume (ESV) [uL] | 278 | 213 |
| Reported End-diastolic pressure (EDP) [mmHg] | 4.49+/-0.51 | 6.69+/-1.10 |
| Model End-diastolic pressure (EDP) [mHg] | 5.42 | 10.9 |

#### 2.2.2 Implementing growth in the compartmental model

We used a kinematic growth framework (Rodriguez et al. 1994) within the compartmental model to simulate LV growth using previously published methods (Witzenburg and Holmes 2018). In this framework, the observed total stretch (*F*_*tot*_) is a product of the growth stretch (*F*_*g*_) and the elastic stretch (*F*_*e*_):

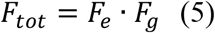

where *F*_*g*_ is a diagonal tensor. The components of *F*_*g*_ were calculated based on the network model CellArea output as described in Section 2.3 below. We modeled the LV as a thin-walled spherical pressure vessel with an initial unloaded cavity volume (*V*_*0*_), radius (*r*_*0*_), and thickness (*h*_*0*_). At each growth step, the growth tensor (*F*_*g*_) was applied to update the unloaded dimensions while assuming no changes in the intrinsic material properties of the myocardium. These dimensions were then used to calculate growth-induced changes in the LV compartmental parameters. After applying growth, multiple heartbeats were simulated until a steady state was reached (compartmental volumes at the beginning and end of the cardiac cycle differed by <0.001ml). Total fiber stretch was calculated throughout the cardiac cycle based on the total chamber dimensions, then divided by the growth stretch to obtain the elastic fiber stretch:

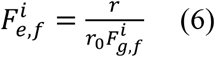

where *r* is the deformed radius of the LV.

### 2.3 Multiscale cardiac growth model

The multiscale cardiac growth model was built by coupling the signaling network model of cell-level growth (Fig. 1, Section 2.1) to the organ-level compartmental growth model (Fig. 2, Section 2.2). Figure 3 provides a schematic that summarizes the coupling between the two models, similar to methods developed in (Estrada et al. 2020). To couple the two models together, we used a linear transfer function to map the elastic stretch calculated by the compartmental model (Eq. 6) to the network Stretch input, and a separate transfer function to map the network CellArea output into a growth stretch (*F*_*g*_). Because the network model employs a single normalized input weight for Stretch between 0 and 1, we chose to map the maximum elastic fiber stretch during a cardiac cycle, max(*F*_*e,f*_). We based this choice on previously published mechanics-based growth models that were successful in predicting growth in response to volume overload using the same mechanical stimulus (Kerckhoffs et al. 2012; Witzenburg and Holmes 2017, 2018). Circulating hormone concentrations (*c*_*AngII*_, *c*_*E2*_, *c*_*P4*_) were mapped to their respective network input weights using separate linear transfer functions:

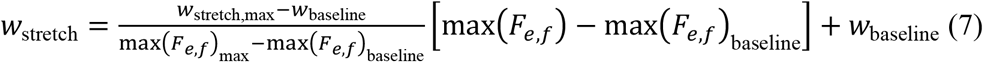

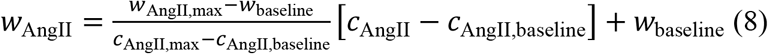

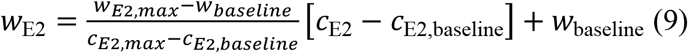

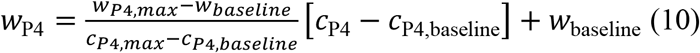

**Figure 3:**
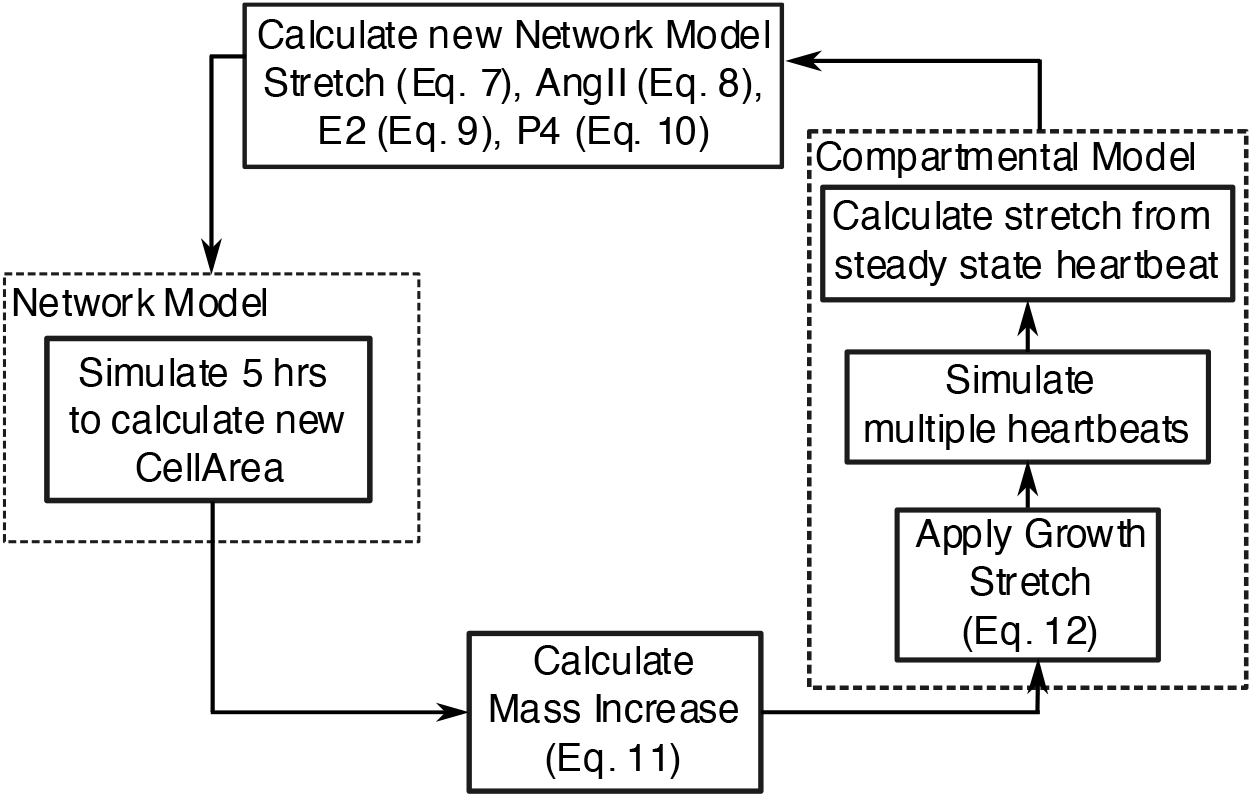
Multiscale model coupling schematic. During each growth step, the compartmental model simulates multiple heartbeats and calculates max(*F*_*e,f*_) experienced by the LV from the steady state heartbeat. We then calculate network model inputs using Eqs. 7-10 and simulate 5 hours of growth to calculate a new CellArea. We then convert CellArea to a growth stretch (*F*_*g*_) using Eqs. 11-12, and apply that growth stretch to the compartmental model before moving to the next growth step.

Here, *w*_*stretch,max*_, *w*_*AngII,max*_, *w*_*E2,max*_, *w*_*P4,max*_ represent the network inputs at which the individual effect of that input on CellArea saturates (Fig. 4A-D). *w*_*baseline*_ is the baseline input weight for each factor. *c*_AngII,baseline_, *c*_E2,baseline_, *c*_P4,baseline_ are the known circulating concentrations of AngII, E2, P4 at the baseline state, and max(*F*_*e,f*_)_baseline_ is the baseline maximum fiber stretch calculated by the compartmental model (see section 2.3.1 below for further details). This leaves one unknown parameter to be optimized for each equation: max(*F*_*e,f*_)_max_, *c*_AngII,max_, *c*_E2,max_, *c*_P4,max_, which represent the elastic fiber stretch and concentrations of AngII, E2, and P4 that trigger maximal remodeling.

**Figure 4:**
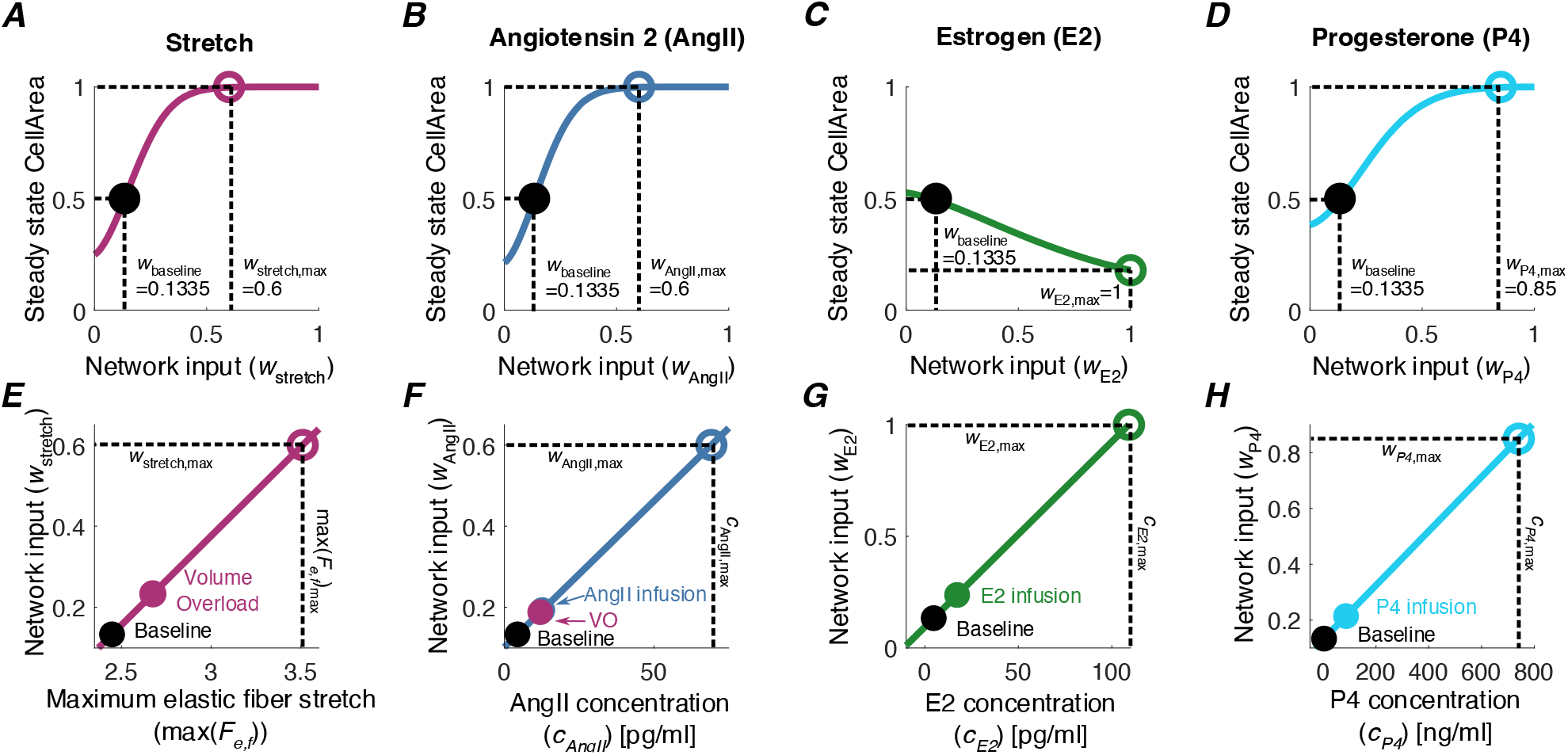
Mapping compartmental model stretch and hormone concentrations to network inputs. Top Row: Curves demonstrate how each network input, A.) Stretch, B.) AngII, C.) E2, and D.) P4 affect the steady state CellArea in the network model. Black dot represents the baseline state, where the input weight, *w*_baseline_ leads to CellArea=0.5. Colored open circles represent the normalized inputs (*w*_stretch,max_, *w*_AngII,max_, *w*_E2,max_, *w*_P4,max_) at which its effect on CellArea saturates. Bottom row, lines represent the linear transfer functions for the network model inputs E.) Stretch, Eq. 7, F.) AngII, Eq. 8, G.) E2, Eq. 9, and H.) P4, Eq. 10. Horizontal axes represent stretch calculated by the compartmental model or hormone concentrations. Black filled circles represent baseline input levels, colored open circles represent the maximum growth response triggered by each stimulus in isolation, and colored filled circles represent the levels of stretch/hormones used to simulate volume overload or hormone infusion.

To translate the growth predicted by network model into a growth stretch for the compartmental model, we used a linear transfer function to map the network output, CellArea, to the determinant of the growth stretch, *J*_*g*_ = det(*F*_*g*_). The parameters for this transfer function were set such that a half-maximal CellArea = 0.5 corresponded to no growth (*J*_*g*_ = 1) and the maximum CellArea = 1 corresponded to a doubling in mass (*J*_*g*_ =2):

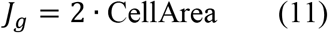

Because heart growth during pregnancy results in both cavity dilation and wall thickening (Savu et al. 2012), we assumed isotropic growth such that the total increase in mass (*J*_*g*_) was evenly distributed in the fiber (*f*), crossfiber (*c*), and radial (*r*) directions. Accordingly, the components of *F*_*g*_ were calculated as:

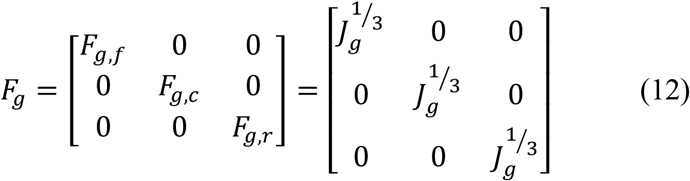

The code for this multiscale model is freely available to download on SimTK (https://simtk.org/projects/heartgrowthpreg).

#### 2.3.1 Calibrating the multiscale model: VO, AngII, E2, P4

To calibrate the multiscale model, we optimized the unknown parameters of the linear transfer functions (Eqs. 7-10, Fig. 4E-H) for Stretch (max(*F*_*e,f*_)_max_), AngII (*c*_AngII,max_), E2 (*c*_E2,max_), and P4 (*c*_P4,max_) to match reported LV growth from studies of experimental VO and infusion of AngII, E2, and P4. In the network model, a normalized input weight of *w*_*Stretch*_ *= w*_*AngII*_ = *w*_*E2*_ = *w*_*P4*_ = 0.1335 produces a steady-state CellArea = 0.5. Thus, max(*F*_*e,f*_)_baseline_ from the baseline compartmental model, *c*_AngII,baseline_ from a control male rat (Ruzicka and Leenen 1995), and *c*_E2,baseline_ (Jankowski et al. 2001) and *c*_P4,baseline_ (Blair and Mickelsen 2006) from an ovariectomized female rat were mapped to *w*_*Stretch*_, *w*_*AngII*_, *w*_*E2*_, and *w*_*P4*_ values of 0.1335 respectively (Supplemental Table 2). Accordingly, the baseline condition in our model represents a nonpregnant ovariectomized female rat or a normal male rat.

To obtain a second point for calibration for each input, we simulated VO and hormonal infusion studies. Since previous studies suggest similar growth between ovariectomized female and male rats (Gardner et al. 2002; Brower et al. 2003) and because studies using ovariectomized female rats are limited, we calibrated our model to growth data from ovariectomized females and male rats with the exception of P4, where only data from non-ovariectomized female mice were available (Chung et al. 2013). For all growth simulations (Table 3), we simulated 21 days of growth based on the rat gestation period.

**Table 3:**
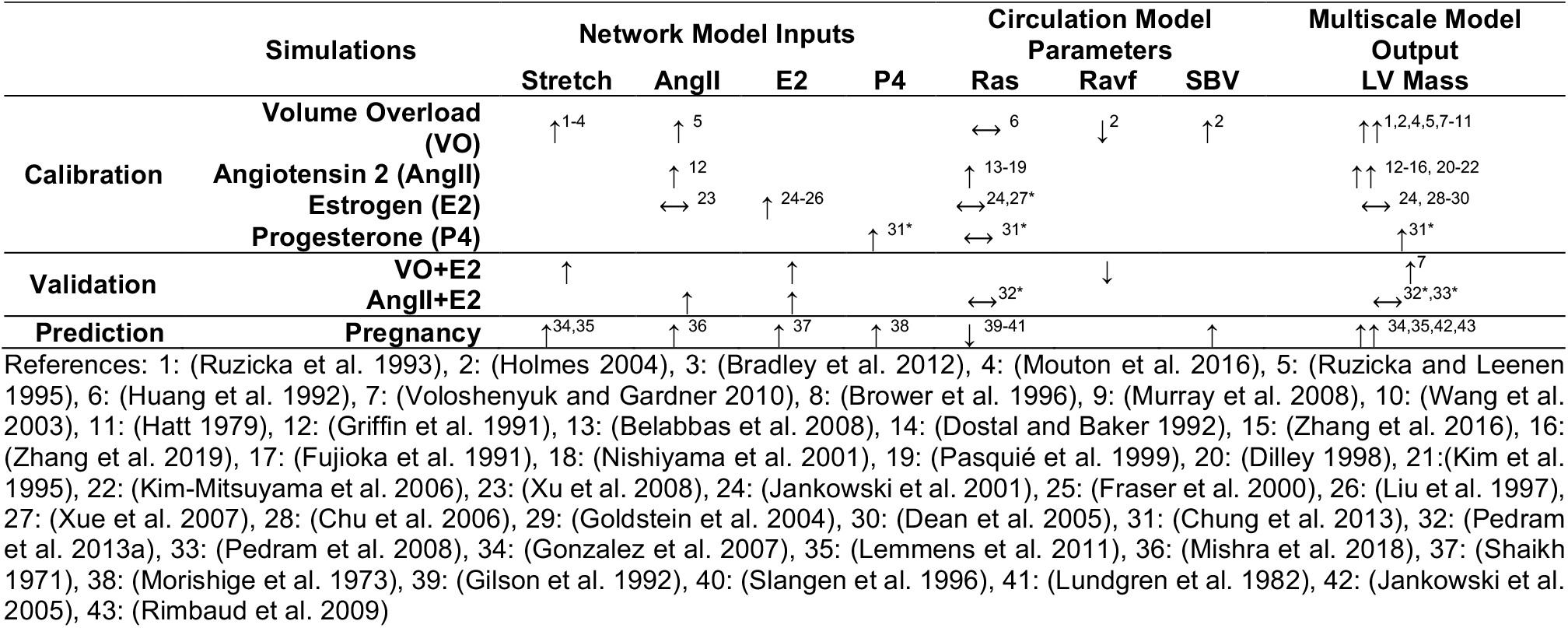
Summary of growth simulations. The multiscale model was calibrated to experimental volume overload (VO) and hormonal infusions of angiotensin 2 (AngII), estrogen (E2), and progesterone (P4) (Section 2.3.1). We then validated model predictions against *in vivo* growth trends from experimental VO+E2 and AngII+E2 (Section 2.3.2). Finally, we predicted heart growth due to pregnancy changes in hormones and circulation model parameters (Section 2.3.3). Arrows represent reported changes from experiments.

Because VO induces changes in AngII levels (Ruzicka and Leenen 1995), we calibrated the linear transfer functions for Stretch (Eq. 7, Fig. 4E) and AngII (Eq. 8, Fig. 4F) simultaneously using reported relative changes in LV mass from VO and AngII infusion experiments in rats. To simulate VO, we set the circulatory parameters to the fitted acute VO values and maintained them throughout growth. Simultaneously, we adjusted *c*_*AngII*_ to match reported values (Ruzicka and Leenen 1995) while keeping baseline *c*_*E2*_ and *c*_*P4*_. To simulate experimental AngII infusion, we induced a step increase in *c*_*AngII*_ while holding baseline *c*_*E2*_ and *c*_*P4*_. In order to capture the fact that *in vivo* AngII infusion gradually increases blood pressure (Fujioka et al. 1991), we linearly increased the arterial resistance (*R*_*as*_) by 3-fold over 21 days of growth.

After calibrating the Stretch and AngII inputs, we simulated LV growth due to E2 and P4 separately to calibrate Eqs. 9 and 10 (Fig. 4G, H). Similar to the AngII simulation, we induced a step increase in *c*_*E2*_ and *c*_*P4*_ while maintaining circulation parameters at baseline. In our review of the literature, we found a wide range in the reported concentrations of E2 and P4. Since no single study reported circulating E2 and P4 concentrations for ovariectomized, E2 or P4 treated, and pregnant animals, we assumed that the infusion experiments induced E2 and P4 concentrations mid-way between nonpregnant and peak pregnant levels. The references for the experimental LV mass we matched for VO, AngII, E2, and P4 infusions are available in Supplemental Data 2: CalibrationGrowth.

#### 2.3.2 Validating the multiscale model: VO+E2 and AngII+E2

After calibrating the multiscale model, we validated its ability to capture interactions between inputs. Because *in vivo* experimental heart growth data were only available for the combinations of VO+E2 (Voloshenyuk and Gardner 2010) and AngII+E2 (Pedram et al. 2013a), we validated our model predictions for these two combinations. To simulate VO+E2, we implemented the same conditions as VO (described above) with an additional step increase in E2. To simulate AngII+E2, we implemented a set increase in AngII and E2, but without an increase in *R*_*as*_, since blood pressures did not increase experimentally in mice treated with AngII+E2 (Pedram et al. 2013a) (Table 3).

#### 2.3.3 Predicting heart growth during pregnancy

After calibrating and validating the multiscale model, we simulated heart growth during pregnancy by incorporating reported changes in hormones and hemodynamics. For the hormonal inputs, we implemented reported changes in *c*_*E2*_ (Shaikh 1971), *c*_*P4*_ (Morishige et al. 1973), and *c*_*AngII*_ (Mishra et al. 2018) from Sprague-Dawley rats during pregnancy: E2 increases slightly in early pregnancy and reaches a peak during late pregnancy, P4 increases in early pregnancy and continues to rise and peaks in late pregnancy before suddenly dropping before delivery, and AngII increases towards the end of pregnancy. Starting in early pregnancy (by gestation day 8 of 21 in rats), cardiac output increases while mean arterial pressure (MAP) remains constant, indicating an overall decrease in systemic resistance (*R*_*as*_). To simulate this decrease in *R*_*as*_, we specified heart rate (*HR*) and calculated changes in *R*_*as*_ from the reported MAP, stroke volume (SV), and *HR* as *R*_*as*_ = MAP/(SV×*HR*) (Gilson et al. 1992; Slangen et al. 1996). Total blood volume also increases during pregnancy due to an increase in both plasma volume and red blood cells (Barron et al. 1984). To simulate this blood volume increase during pregnancy (Pritchard 1965; Barron et al. 1984), we allowed *SBV* to change at each growth step to match the reported time course of end-diastolic diameter (EDD) during pregnancy (Table 3).

#### 2.3.4 Uncertainty analysis

Finally, to understand how variability in experimental measurements of the model inputs affect its predictions, we conducted uncertainty analyses for the two primary drivers of growth in the model during pregnancy, P4 and hemodynamics. First, to test how uncertainty in P4 measurements affect model predictions, we simulated a high P4 time course by implementing hormone levels at mean plus one standard deviation of the experimental data, and a low time course of P4 as the experimental mean minus 1 SD. Similarly, to test how uncertainty in hemodynamic measurements affect predictions, we allowed for *SBV* compensation at every growth step to match high or low EDD time course.

## 3. Results

### 3.1 Calibrating the model: Multiscale model captures both growth and hemodynamics due to VO, AngII, E2, and P4

We were able to quantitatively capture the reported relative changes in LV mass due to experimental VO and infusion of the hormones, AngII, E2, and P4 by optimizing the linear transfer function parameters in Equations (7)-(10) (Fig. 5A, Supplemental Table 2). We confirmed that the optimized parameters minimized error between model predictions and experimental growth data by conducting a sensitivity analysis (Supplemental Figure 2). Experimental VO and infusion with AngII in rats led to similar amounts of growth at 21 days, with LV mass increasing 30% for VO and 25% for AngII. P4 infusion led to less growth, increasing LV mass by 13%, which was reported to be significant in the experiment (Chung et al. 2013). E2 infusion led to a small, non-significant atrophy with a 5% decrease in LV mass. In addition to matching growth data, the specified circulatory parameters (Table 3) resulted in good agreement with the reported hemodynamic data for early VO and AngII infusion, including a step increase in cardiac output (CO) with VO and a gradual decrease in CO with AngII infusion (Fig. 5B). No changes in CO occurred in the E2 and P4 simulations, which agreed with previously reported experimental observations in mice (Babiker et al. 2006; Chung et al. 2013). Real-world and calibrated network inputs for hormone levels and stretch throughout these simulations are shown in Figure 6. Notably, the Stretch input (Fig. 6D) increased acutely and then gradually decreased in our VO simulations while it remained relatively constant near baseline for AngII, E2, and P4.

**Figure 5:**
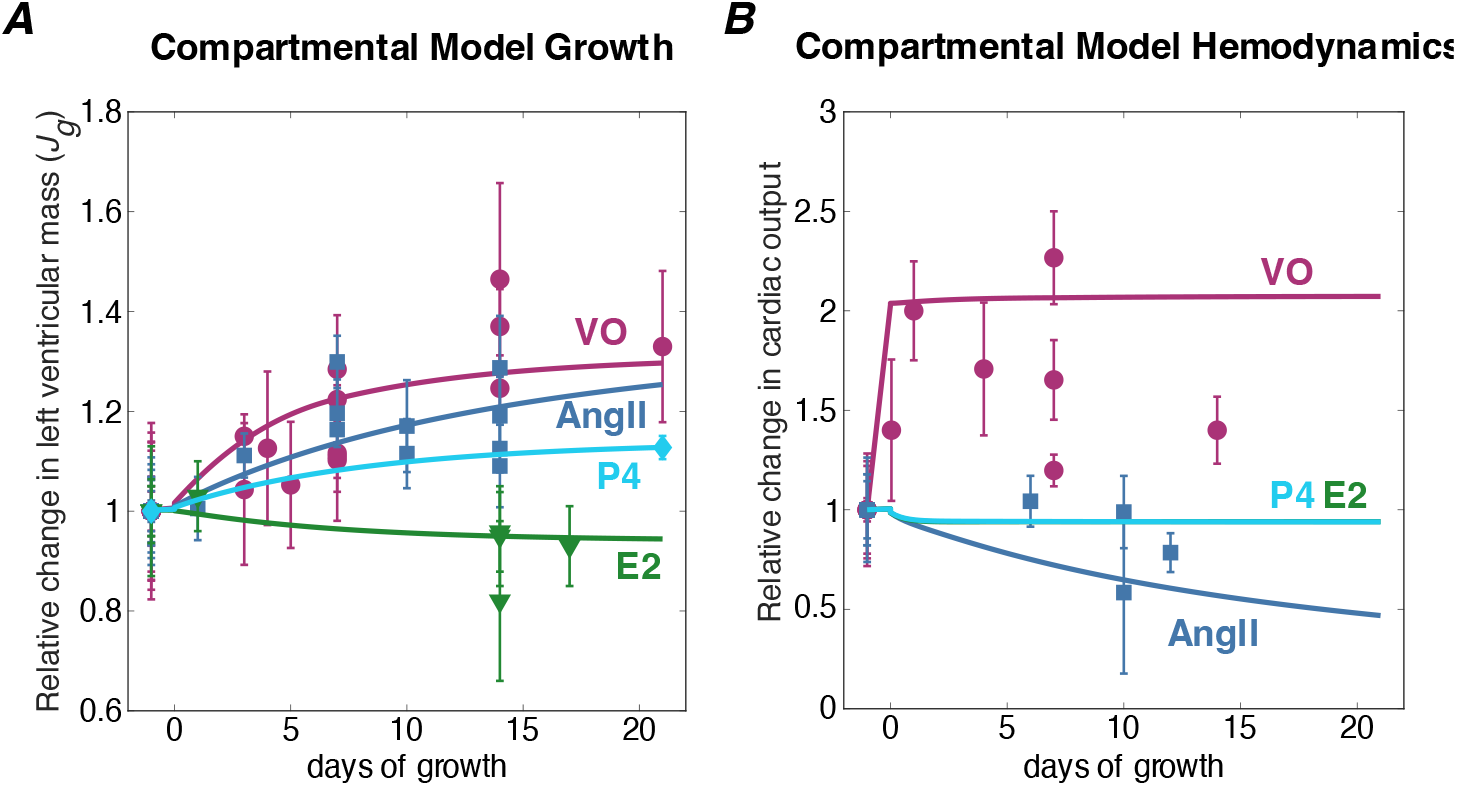
Calibrating the multiscale model. A.) Model match to experimental data for each experimental condition. Growth quantified as relative change in left ventricular mass and B.) Hemodynamics quantified as relative change in cardiac output for VO, AngII, E2, and P4. Symbols represent reported data +/-1 standard deviation (see Table 3) and lines represent model outputs. More details including references for the experimental studies are available in Supplemental Data 2: CalibrationGrowth. (Online version in color).

**Figure 6:**
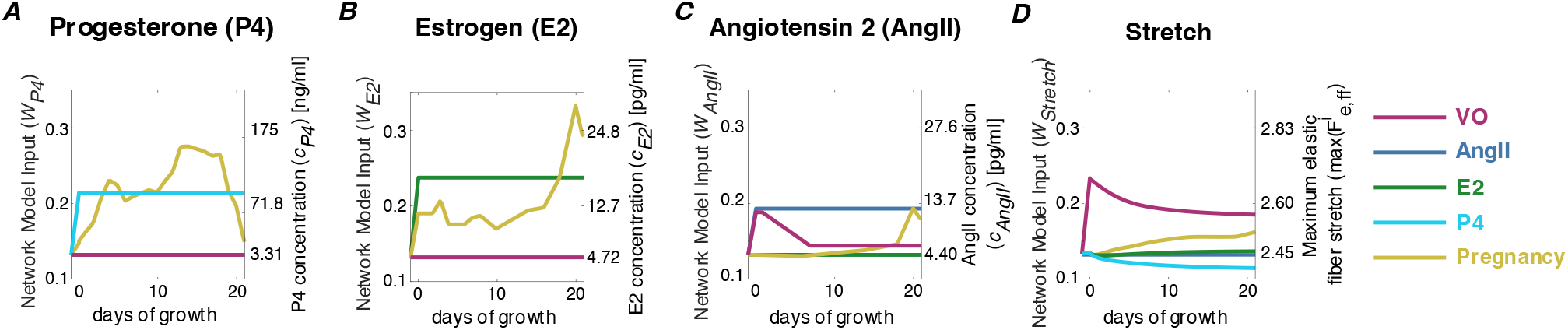
Calibrated hormone and stretch inputs for the network model during simulations of volume overload (VO, purple), Angiotensin 2 (AngII, blue) infusion, Estrogen (E2, green) infusion, and Progesterone (P4, cyan) infusion. Gold lines represent input levels during pregnancy simulation (Section 3.3). Right axes show hormone concentrations in (pg/mL) or (ng/mL) and maximum elastic fiber stretch from the compartmental model. Left axes show calibrated, normalized network inputs calculated from Equations (7)-(10). (Online version in color).

### 3.2 Multiscale model correctly predicts attenuated growth trends with VO+E2 and AngII+E2

When we simulated the combinations of VO+E2 and AngII+E2, the model correctly predicted the trends in attenuated growth compared to VO and AngII alone. Our VO+E2 simulation led to 4% attenuation in LV mass compared to VO alone (Fig. 7A). These results agree with experiments where ovariectomized rats with VO+E2 exhibited slight, insignificant attenuation in growth with reported 5% LV mass attenuation at 5 days and 18% attenuation at 8 weeks (Voloshenyuk and Gardner 2010) compared to ovariectomized rats subjected to VO alone. We note that the VO+E2 study employed VO of much longer duration of 56 days compared to our calibration studies and exhibited more growth, which could explain why our model underestimated the amount of attenuation.

**Figure 7:**
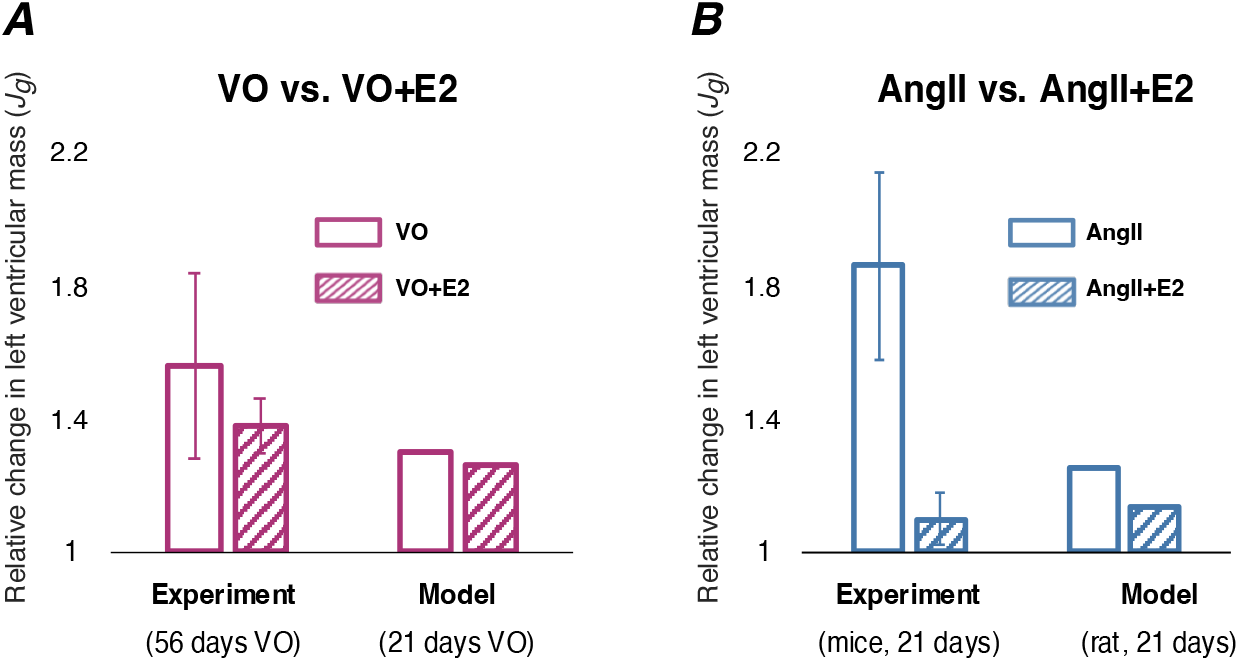
Validating the multiscale model. Simulations of VO+E2 and AngII+E2 (shown in dashed lines) both led to attenuated growth compared to VO only and AngII only. A.) Comparisons between experimental VO and VO+E2 at 56 days in rats (Voloshenyuk and Gardner 2010) versus model (21 days of VO). B.) Comparisons between experimental AngII and AngII+E2 at 21 days in mice (Pedram et al. 2013a) versus model (21 days of simulated AngII infusion in rats). Error bars represent 1 standard deviation.

Our simulations of AngII+E2 led to more attenuation (12% attenuation in LV mass compared to AngII alone) compared to our simulations of VO+E2 (Fig. 7B). Ovariectomized mice treated with AngII+E2 exhibited significant attenuation in growth with reported 65% LV mass attenuation compared to AngII alone after 21 days of treatment (Pedram et al. 2013a). Again, the model underestimated the amount of attenuation, which could be due to the use of mice in the experiments where the model simulates a rat anatomy. Our network model in these *in vivio* simulations led to similar activations for CaN, Akt, and GSK3B as our *in vitro* simulations (Fig. 1B, AngII+E2 column). This result was in agreement with the reported similar activity of CaN, Akt, and GSK3B between *in vitro* cell culture experiments and *in vivo* animal experiments (Pedram et al. 2008, 2013a).

### 3.3 Multiscale model predicts realistic changes in growth and hemodynamics during pregnancy

When we incorporated hormones and circulation parameter changes representative of pregnancy, our calibrated and validated multiscale model predicted growth and hemodynamic changes that were in good agreement with the experimental data. Over 21 days of pregnancy, LV mass gradually increased by approximately 19% (Fig. 8A). These model predictions were within the reported data range, except during early pregnancy where the model slightly overpredicted growth. By allowing for *SBV* compensation to match the reported change in end-diastolic diameter (Fig. 8B) (Gonzalez et al. 2007; Lemmens et al. 2011), the model predicted changes in MAP (Fig. 8C) and CO (Fig. 8D) that were in good agreement with the reported data (Gilson et al. 1992; Slangen et al. 1996; Linke et al. 2002). Overall, our model predicted a 29% increase in *SBV* at day 14 of pregnancy before decreasing slightly towards the end of pregnancy.

**Figure 8:**
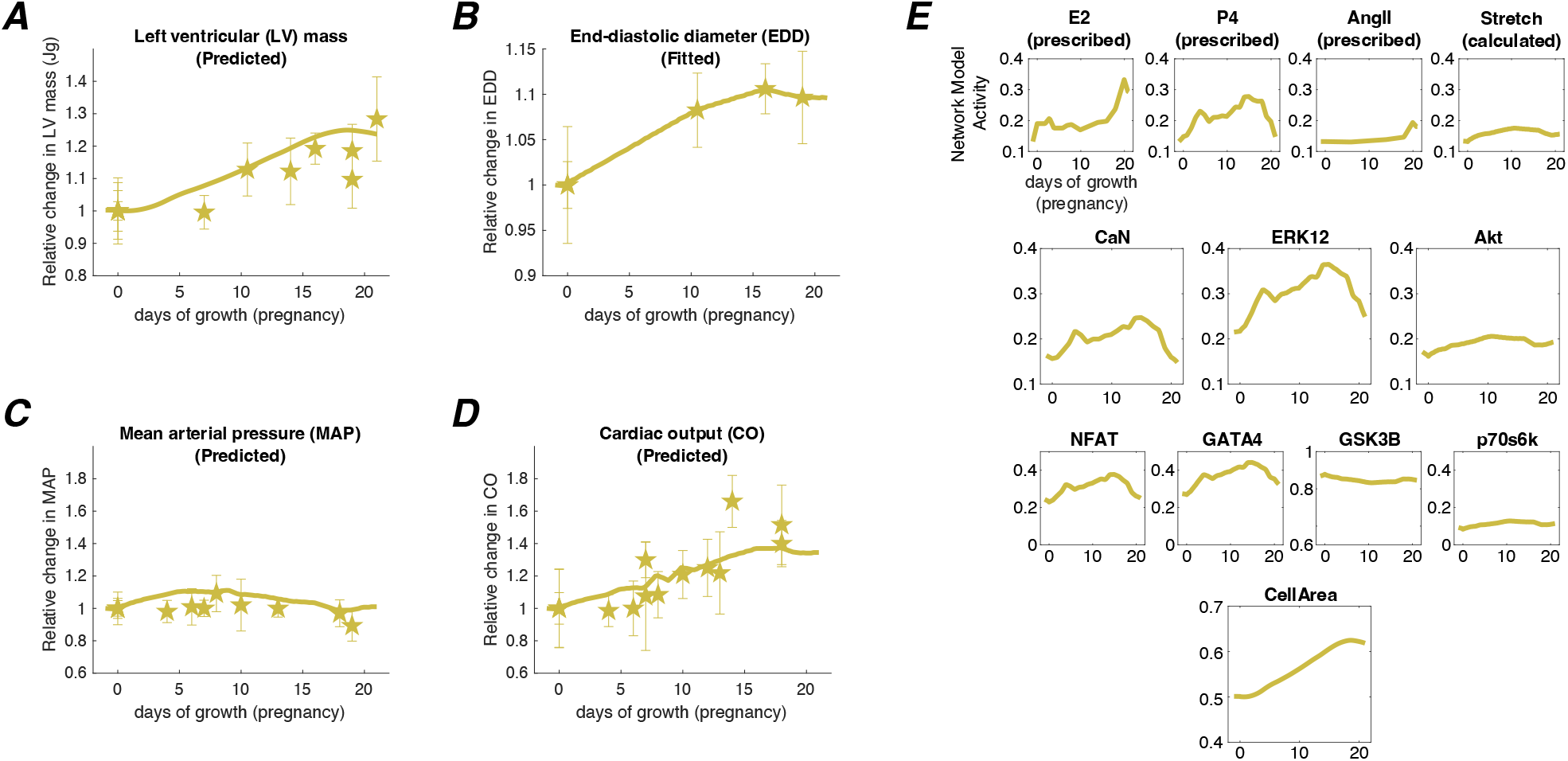
Multiscale model predictions for heart growth and hemodynamic changes during pregnancy. A.) Relative change in left ventricular mass, B.) Relative change in end-diastolic cavity diameter (EDD), C.) Relative change in mean arterial pressure (MAP), and D.) Relative change in mean cardiac output (CO). Symbols represent experimental mean +/-1 standard deviation and solid lines represent model predictions. References for experimental data are available in Supplemental Data 2: PregnancyGrowth. E.) Activity of nodes in network model during pregnancy simulations. Each panel shows changes in normalized model weights that range from 0 to 1 for one node in the signaling network (Figure 1A).

An advantage of this multiscale modeling approach is the ability to examine intracellular signals that drive the organ-level growth. When we examined the predicted network response for the pregnancy simulation, cell-level growth coincided with network species that are downstream of P4 (CaN, ERK12, NFAT, GATA4) (Fig. 8E). Surprisingly, the network Stretch input did not change significantly, and there was minimal activation of stretch-related signaling pathways (Akt, GSK3B, p70s6k). In our model, the elastic stretch, and therefore the network Stretch input, is dependent on the rate at which the LV volume increases versus the rate at which the ventricle grows. Because the LV growth driven by high P4 levels in early pregnancy (Fig. 8E) balanced the change in LV volume/EDD (Fig. 8B), our model predicted minimal change in the elastic stretches. To further understand the relative contributions of P4 and Stretch on growth during pregnancy, we ran additional pregnancy simulations where we maintained either P4 or Stretch at baseline (*w*_P4_ = 0.1335 or *w*_Stretch_ = 0.1335) and prescribed the same *SBV* changes as in the original pregnancy simulations without attempting to match the reported EDD. Both cases underpredicted growth to similar degrees (Fig. 9A). Together, these results suggest the important roles of both P4 and hemodynamics in driving pregnancy-induced heart growth.

**Figure 9:**
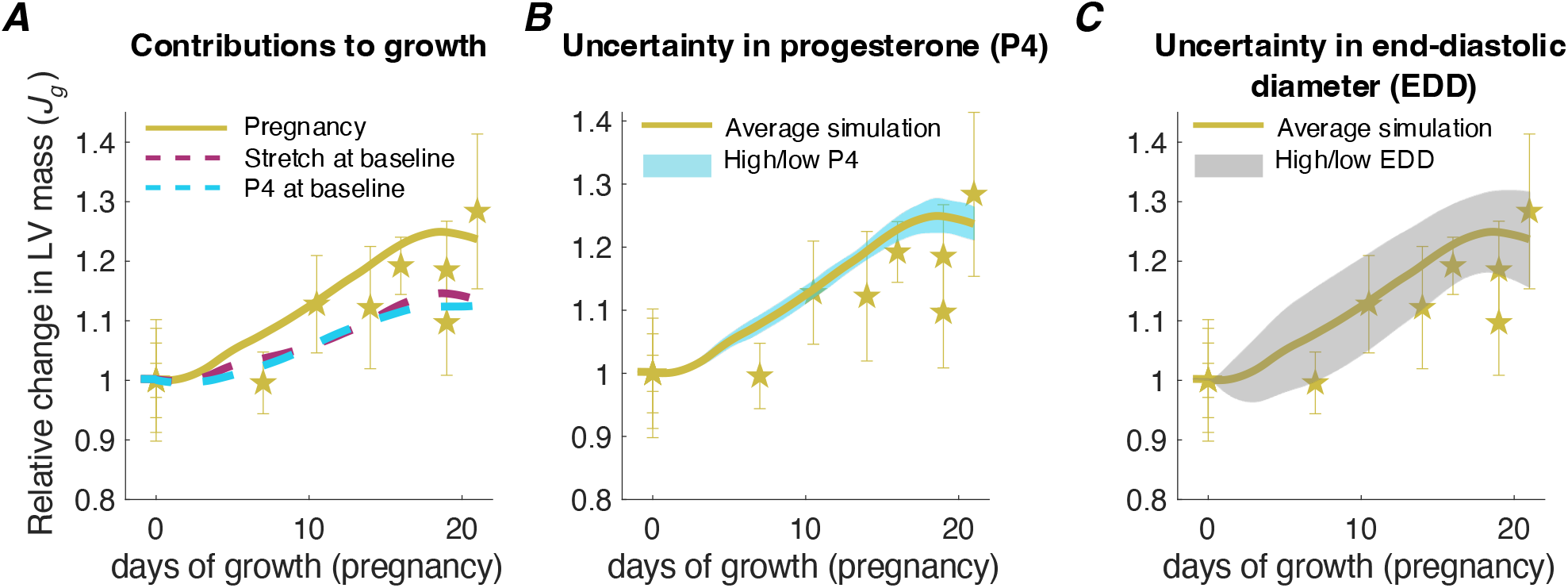
Effects of progesterone (P4) and hemodynamics on heart growth during pregnancy. A.) To understand the contributions of P4 and Stretch to LV growth, pregnancy simulations were repeated while maintaining P4 at baseline (dashed blue line) or Stretch at baseline (dashed purple line). To understand how the reported uncertainty in P4 and hemodynamics affect growth predictions, simulations were repeated with high (mean + 1 standard deviation) and low (mean - 1 standard deviation) levels of B.) P4 and C.) end-diastolic diameter. Symbols represent experimental mean +/-1 standard deviation. Range of model predictions are represented by the shaded areas. Solid gold line represents the original pregnancy simulation with average values.

### 3.4 Variability in P4 and hemodynamics translate to appropriate variability in predicted growth

Finally, to investigate how uncertainty in the experimental measurements that determine inputs to our model would influence its growth predictions, we conducted additional simulations with different levels of progesterone and hemodynamic loading. When we repeated our pregnancy simulations with high (mean plus 1 standard deviation of the reported data) and low (mean minus 1 standard deviation of the reported data) levels of P4 (Morishige et al. 1973), the multiscale model predicted a +/-2% LV mass change compared to our original simulations (blue area, Fig. 9B), which was smaller than the standard deviation in reported LV mass at most time points. We also repeated our pregnancy simulations while adjusting the *SBV* to match high and low levels of the reported EDD (mean plus or minus 1 standard deviation of the reported data), which produced +/-8% variations in LV mass change, similar to the reported standard deviations of LV mass change (gray area, Fig. 9C). Together, these results suggest that our calibrated multiscale model provides growth predictions consistent with experimental data, with an appropriate sensitivity to known variations in the factors that drive hypertrophy during pregnancy.

## 4 Discussion

The objective of this work was to develop a multiscale cardiac growth model that incorporates interactions between mechanical and hormonal signaling contributing to heart growth during pregnancy. To build this model, we coupled an intracellular signaling network model of cell-level growth to a compartmental mechanical model of organ-level growth. We first calibrated this multiscale model to heart growth in response to experimental VO and infusion experiments of AngII, E2, and P4. We then validated the model’s ability to capture interactions between inputs by confirming predictions of attenuated heart growth for the combinations of VO+E2 and AngII+E2 as reported in the literature. When we simulated pregnancy by introducing appropriate changes in hormonal inputs and hemodynamics to our calibrated model, we were able to produce growth predictions that were in good agreement with experimental data. Furthermore, simulating the reported variability in progesterone concentrations and hemodynamic measures translated to appropriate variability in the predicted growth. Interestingly, the rise in progesterone that occurs during early pregnancy drove LV growth in our model, resulting in minimal changes in the elastic stretch in the organ-level model, and thus minimal stretch-activated signaling in the network model. This result suggests that progesterone stimulation might be sufficient to account for much of pregnancy-induced heart growth in the rat, particularly during the early stages of pregnancy.

In our model, changes in the elastic stretch, and therefore network model Stretch, are dependent on the rate at which LV end-diastolic volume (EDV) increases compared to the rate at which the ventricle grows. During experimental VO, a step increase in EDV occurs immediately as end-diastolic pressure (EDP) and cardiac output rise following creation of the overload (Table 2). Therefore, the rate of LV EDV increase clearly outpaces LV growth and the network component of the model experiences and responds to increased Stretch (Fig. 6D, VO simulation). Conversely, in our pregnancy simulation, the rise in progesterone that occurs in early pregnancy leads to growth that approximately matches the increase in EDV as *SBV* gradually increases throughout pregnancy. Thus, elastic stretch in the compartmental model and myocyte Stretch in the network model remain near baseline levels. This prediction, however, is difficult to verify since elastic stretch is a theoretical construct that cannot be directly measured. One approach to explore this prediction further might be to fix hearts at their *in vivo* end-diastolic diameters and measure sarcomere lengths as an indicator of cell-level stretch. Alternatively, stretch-induced signaling activity could be assayed; however, as discussed below the available data on stretch-induced signaling during pregnancy are too variable to support any firm conclusions.

A novel advantage of the modeling framework presented here is the ability to validate our predictions across multiple length scales, from intracellular signaling to organ-level growth. When we examined the predicted network response in our pregnancy simulations, species downstream of P4 (CaN, ERK12, NFAT, GATA4) increased in early pregnancy and peaked in late pregnancy, before dropping back to baseline levels by the end of the gestation (Fig. 8E). These trends generally align with the reported levels of these species collected from mice and rat hearts throughout pregnancy (Gonzalez et al. 2007; Lemmens et al. 2011; Chung et al. 2012, 2013; Xu et al. 2016). However, the minimal changes in the signaling intermediates downstream of Stretch (Akt, GSK3B, p70s6k) (Fig. 8E) disagree with some published data. With respect to Akt, some studies have reported an increase during pregnancy in mice and rats (Hilfiker-Kleiner et al. 2007; Lemmens et al. 2011; Chung et al. 2012), one study reported a decrease towards late pregnancy in rats (Gonzalez et al. 2007), and another study reported no change during late pregnancy in mice (Xu et al. 2016). GSK3B and p70s6k have been reported to rise during mid-pregnancy before decreasing to nonpregnant levels by late gestation in mice (Chung et al. 2012). In one *in vitro* study, P4 did not affect Akt phosphorylation (Chung et al. 2012), while the effect of P4 on GSK3B and p70s6k activity are unknown. To improve our predictions at the network level, there is a need for more experimental data. In particular, a better understanding of how P4 and Stretch interact is necessary. For example, an *in vivo* experiment where VO is induced along with P4 infusion could provide valuable insights into how Stretch and P4 interact to affect heart growth as well as intracellular signaling through Akt, GSK3B, and p70s6k.

In terms of the clinical applicability of this model, there are both similarities and differences between rat and human pregnancy. Generally, hemodynamic changes are similar between rats (Gilson et al. 1992; Slangen et al. 1996; Blair and Mickelsen 2006) and humans (Hunter and Robson 1992) (increase in CO, decrease in systemic resistance, and an increase in blood volume). In terms of heart growth, similar amounts of growth have been reported for rats and human pregnancies (Savu et al. 2012), both in terms of the magnitude and patterns of growth. Hormonal profiles during pregnancy, however, differ slightly between rats and humans in that pregnant women do not exhibit a drop in progesterone before delivery (Tulchinsky et al. 1972). Finally, another big difference between rat and human pregnancy is the average number of offsprings per pregnancy. Rats generally give birth to an average litter of 11-12 pups, while most human pregnancies are singleton pregnancies. Interestingly, CO is known to be higher in patients carrying twins (Hunter and Robson 1992). Before translating this model for human pregnancy, these differences should be further investigated and accounted for.

One limitation of this work is the simplified network model, where we represent a limited number of the many intracellular pathways that can lead to cardiomyocyte growth. As mentioned previously, we chose this simplified network model because experimental data on the effects of E2 and P4 on hypertrophic pathways are currently limited. As more data on how pregnancy hormones affect intracellular hypertrophic pathways become available, an exciting future direction would be to incorporate E2 and P4 into a more comprehensive network model of hypertrophy. The use of a simplified network may also explain why our model captured the trend of attenuated hypertrophy with E2 administration in our validation studies, but underpredicted its magnitude (Fig. 7). Here again, more data are needed to allow apples-to-apples comparisons, since the studies we found that administered E2 during VO or AngII differed significantly from other studies used to tune the model in their degree of baseline hypertrophy in the absence of E2. Another limitation of this model is that we assumed isotropic growth without considering directional growth and shape changes (myocyte thickening leading to wall thickening versus myocyte lengthening leading to cavity dilation). Since pregnancy is associated with isotropic growth and because signaling pathways and mechanical signals that control these patterns of myocyte hypertrophy are currently unknown (Maillet et al. 2013), we focused on matching LV mass change for our simulations. Therefore, this model is unable to predict anisotropic cardiac growth as observed in cases of pressure or volume overload. As the signaling mechanisms that control patterns of hypertrophy become clearer, the next iteration of the model could be extended to distinguish between myocyte thickening and lengthening. Finally, when simulating hemodynamic changes during pregnancy, we implemented the reported changes in systemic resistance without accounting for the direct effects of E2 and P4 on the aorta and vessels. Additionally, we chose to adjust *SBV* to match the reported EDD (a surrogate for EDV). While our simulations produced values of MAP and CO within the reported experimental means, additional experimental data – including simultaneous measurements of LV mass, EDD or EDV, and end-diastolic pressure (EDP) – would provide more confidence in these predictions. While *SBV* is extremely difficult to measure directly, we have found in prior work that fitting EDP allows good estimation of *SBV* changes during multiple different types of overload (Witzenburg and Holmes 2018).

In summary, we present here a computational model of maternal heart growth during pregnancy that considers both hormonal and mechanical stimuli. In this modeling study, we calibrated the model to contributions of hormones and stretch on heart growth and demonstrated its ability to predict realistic cardiac hypertrophy during pregnancy. Furthermore, our analysis suggested that the early rise in progesterone that occurs during pregnancy may account for much of the observed heart growth in rats, particularly during the first half of gestation. Based on our model analysis, we conclude with suggestions for future experimental studies, including more cell-levels studies on the effects of pregnancy hormones on hypertrophic pathways, as well as potential *in vivo* experiments to determine the synergistic effects of elevated stretch and progesterone on heart growth.

## Funding

This work was supported by the National Institutes of Health (U01 HL127654 and R01 HL137755).

## Availability of data and material

Available in the Online Supplement

Code availability: All code developed for this study is available on SimTK (https://simtk.org/projects/heartgrowthpreg); the network model can be run independently through Netflux (https://github.com/saucermanlab/Netflux).

## Compliance with ethical standards

### Conflict of interest

None to declare

### Ethical approval/Consent to participate/Consent for publication

Not applicable.

## Notes

https://simtk.org/projects/heartgrowthpreg

### Competing Interest Statement

The authors have declared no competing interest.

### Summary of Updates

This revision contains a link for the code presented here on SimTK, as well as new sensitivity studies and Monte Carlo simulations to determine the uniqueness of model parameter fits.

