## Supplemental Document for "Multiscale model of heart growth during pregnancy: Integrating mechanical and hormonal signaling"

**Online Supplement**

**Methods**

**1. Complete list of network model equations**

$$\frac{d\text{AngII}}{dt} = \frac{1}{\tau_{\text{AngII}}}[Y_{MAX}(\text{AngII}) - \text{AngII}]$$

$$\frac{d\text{E2}}{dt} = \frac{1}{\tau_{\text{E2}}}[Y_{MAX}(\text{E2}) - \text{E2}]$$

$$\frac{d\text{P4}}{dt} = \frac{1}{\tau_{\text{P4}}}\{Y_{MAX}(\text{P4}) - \text{P4}\}$$

$$\frac{d\text{Stretch}}{dt} = \frac{1}{\tau_{\text{Stretch}}}\{Y_{MAX}(\text{Stretch}) - \text{Stretch}\}$$

$$\frac{d\text{CaN}}{dt} = \frac{1}{\tau_{\text{CaN}}}\{OR[AND(f_{inhib}(\text{E2}), f_{act}(\text{P4})), AND(f_{act}(\text{AngII}), f_{inhib}(\text{E2}))] \cdot Y_{MAX}(\text{CaN}) - \text{CaN}\}$$

$$\frac{dMEK1}{dt} = \frac{1}{\tau_{MEK1}}\left\{OR\left[f_{act}(Stretch), OR\left(AND(f_{act}(AngII), f_{inhib}(E2)), f_{act}(P4)\right)\right] \cdot Y_{MAX}(MEK1) - MEK1\right\}$$

$$\frac{d\text{Akt}}{dt} = \frac{1}{\tau_{\text{Akt}}}\{OR[AND(f_{act}(\text{AngII}), f_{inhib}(\text{E2})), f_{act}(\text{Stretch})] \cdot Y_{MAX}(\text{Akt}) - \text{Akt}\}$$

$$\frac{d\text{ERK12}}{dt} = \frac{1}{\tau_{ERK12}}\{f_{act}(\text{MEK1}) \cdot Y_{MAX}(\text{ERK12}) - \text{ERK12}\}$$

$$\frac{d\text{GSK3B}}{dt} = \frac{1}{\tau_{\text{GSK3B}}}\{f_{inhib}(\text{Akt}) \cdot Y_{MAX}(\text{GSK3B}) - \text{GSK3B})$$

$$\frac{d\text{mTor}}{dt} = \frac{1}{\tau_{\text{mTor}}}\{f_{act}(\text{Akt}) - Y_{MAX}(\text{mTor})\}$$

$$\frac{d\text{NFAT}}{dt} = \frac{1}{\tau_{\text{NFAT}}}\{OR[f_{act}(\text{CaN}), OR[AND(f_{act}(CaN), f_{act}(ERK12)), f_{inhib}(GSK3B)]] \cdot Y_{MAX}(\text{NFAT}) - \text{NFAT}\}$$

$$\frac{d\text{GATA4}}{dt} = \frac{1}{\tau_{\text{GATA4}}}\{\text{OR}[f_{inhib}(\text{GSK3B}), f_{act}(\text{ERK12})] \cdot Y_{MAX}(\text{GATA4}) - \text{GATA4}\}$$

$$\frac{dp70s6k}{dt} = \frac{1}{\tau_{p70s6k}}\{f_{act}(mTor) \cdot Y_{MAX}(\text{p70s6k}) - \text{p70s6k}\}$$

$$\frac{d\text{CellArea}}{dt} = \frac{1}{\tau_{\text{CellArea}}}\left\{OR\left[f_{act}(\text{NFAT}), OR\left(f_{inhib}(\text{GSK3B}), OR(f_{act}(\text{GATA4}), f_{act}(\text{p70s6k}))\right)\right] \cdot Y_{MAX}(\text{CellArea}) - \text{CellArea}\right\}$$

**2. Allometric Scaling for Circulation Model Parameters**

We utilized allometric scaling (West et al. 1997) to scale the circulation model from a canine to a rat anatomy and circulation. We based the canine circulation values on our groups previous model (Witzenburg and Holmes 2018). The allometric scaling law assumes that a biological variable for a rat, $Y_{rat}$, is dependent on the same biological variable for a canine, $Y_{canine}$, and the body mass ratio of rat to dog, $M_{rat}/M_{canine}$, through the following relationship:

$$Y_{rat} = Y_{canine}\left(\frac{M_{rat}}{M_{canine}}\right)^{b}$$

Here, b is the scaling exponent for a specific biological variable. For this study, we used b = -3/4 for resistors and b = 1 for capacitors in the circulation model as reported (West et al. 1997) and assumed an average canine weight of 20 kg (Santamore and Burkhoff 1991) and an average rat weight of 260 g (Slangen et al. 1996; Holmes 2004; Jankowski et al. 2005; Gonzalez et al. 2007; Lemmens et al. 2011). Accordingly, we calculated the following model parameters for the rat model (Supplemental Table 1). Here, we fit the resistances $R_{as}$ and $R_{cs}$ against the available experimental data (See main manuscript, Section 2.2.1 for more details).

**Supplemental Table 1:** Circulation parameters for the Canine model (as published in (Witzenburg and Holmes 2018) and the calculated circulation parameters for the rat model for the current study using allometric scaling.

| | **Canine (Witzenburg, 2018), $M_{canine}$ = 20,000g** | **Rat (Current study), $M_{rat}$ = 260g** |
|---|---|---|
| **Resistances (b=-3/4)** | | |
| Pulmonary venous resistance $R_{vp}$ | 0.015 | 0.39 |
| Characteristic resistance of the aorta $R_{cs}$ | 0.023 | Fitted against data |
| Systemic arterial resistance $R_{as}$ | 1.6 | Fitted against data |
| Systemic venous resistance $R_{vs}$ | 0.015 | 0.39 |
| Characteristic resistance of the pulmonary artery $R_{cp}$ | 0.060 | 1.6 |
| Pulmonary arterial resistance $R_{ap}$ | 0.30 | 7.8 |
| **Capacitors (b=1)** | | |
| Pulmonary venous compliance $C_{vp}$ | 0.060 | 0.039 |
| Systemic arterial compliance $C_{as}$ | 0.019 | 0.013 |
| Systemic venous compliance $C_{vs}$ | 0.34 | 0.22 |
| Pulmonary arterial compliance $C_{ap}$ | 0.040 | 0.026 |

### 3. Determining model parameter uniqueness for baseline and acute volume overload hemodynamics

We conducted a Monte Carlo simulation to test whether our optimization for the baseline and acute hemodynamics (Section 2.2.1 in the main manuscript) resulted in a unique set of circulation parameters. We tested 5000 random numerical combinations of the five fitted parameters ($R_{as}$, $R_{cs}$, $R_{avf}$, $SBV_{control}$, $SBV_{VO}$, see Table 2 in the main manuscript), simulated baseline and acute volume overload (VO) pressure-volume behavior, and calculated the error between the predicted model and the reported hemodynamic measures (Holmes 2004). In the first set of 5000 simulations, we tested a range of parameter values that were 50% to 150% of the optimized value (left column, Suppl. Fig. 1). In the second set of 5000 simulations, we tested a range of 95%-105% of the optimized value (right column, Suppl. Fig. 1). Deviations from the optimized parameter combination led to significantly higher errors, suggesting that the optimized parameters from our fits are unique and give the best fit to the experimental data.

<table>
<tr><th></th><th>50-150% of optimized value</th><th>95-105% of optimized value</th></tr>
<tr><td>R<sub>as</sub></td><td>
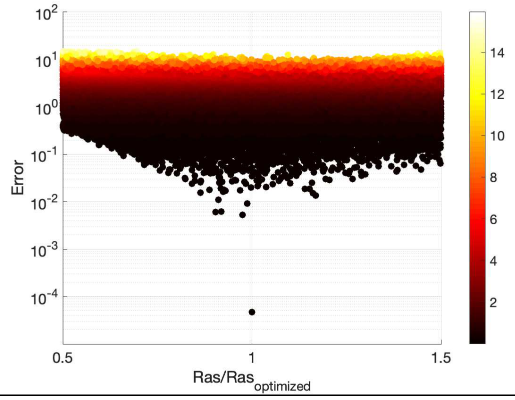

</td><td>
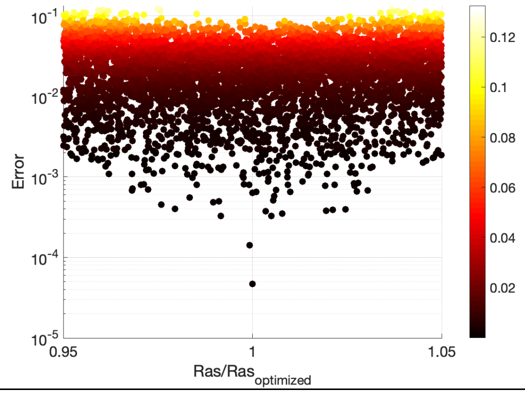

</td></tr>
<tr><td>R<sub>cs</sub></td><td>
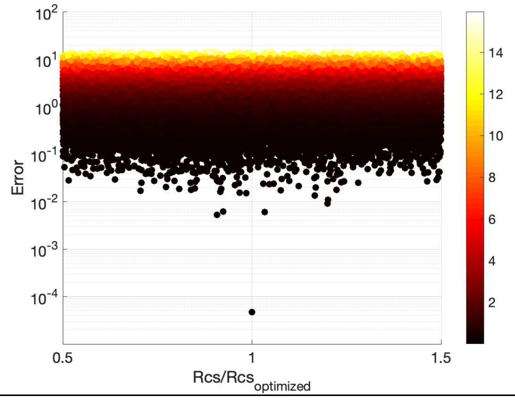

</td><td>
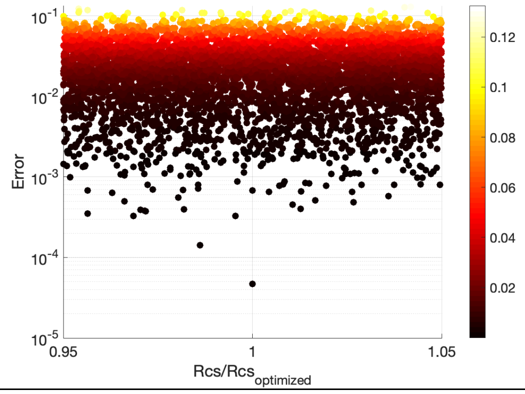

</td></tr>
<tr><td>R<sub>avf</sub></td><td>
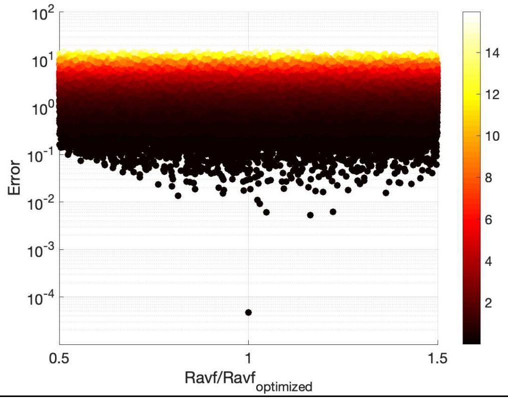

</td><td>
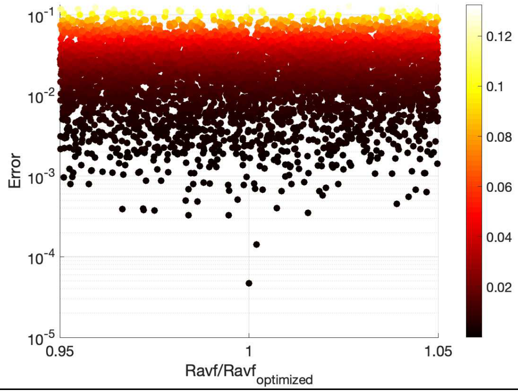

</td></tr>
<tr><td>SBV, control</td><td>
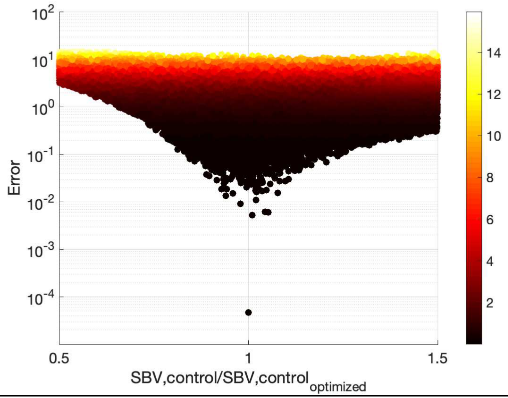

</td><td>
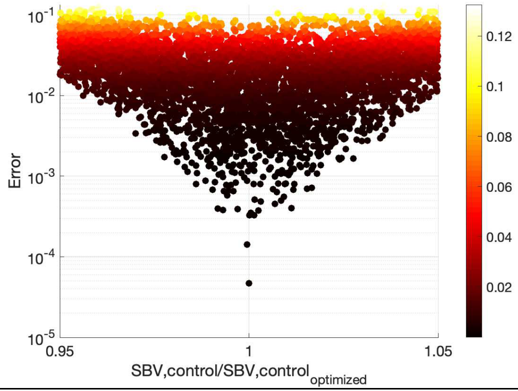

</td></tr>
</table>

**SBV, VO**

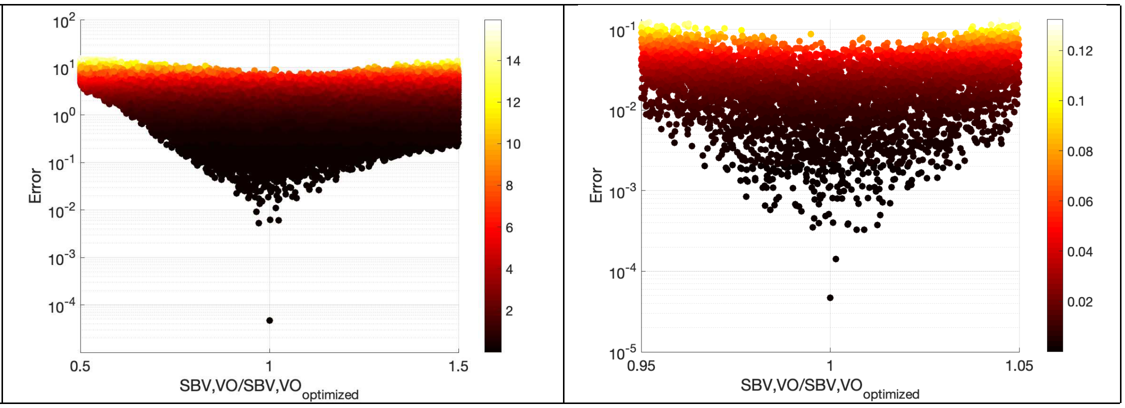


**Supplemental Figure 1: Monte Carlo simulations to determine optimized circulation model parameter uniqueness.** 5000 random numerical combinations of the five fitted circulation parameters (each row) were tested. Left column represents 5000 combinations of parameter values between 50% to 150% of the optimized value. Right column represents 5000 combinations of parameter values between 95%-105% of the optimized value. The horizontal axes show the ratio of the tested circulation parameter to the optimized parameter. The vertical axes represent the calculated error (model against reported hemodynamic data) on a log scale.

**Results:**

**Supplemental Table 2: Linear transfer function parameters for network model inputs.** Baseline network weight was determined as the weight that produces a steady state CellArea =0.5. Maximum network weights are input weights at which no further increases in weights lead to changes in CellArea (see Fig. 3A-D, main manuscript). Baseline stretch/hormone concentrations are mapped to $w_{\text{baseline}}$ and are either calculated by the compartmental model for stretch or are reported hormone levels in normal male or ovariectomized rats. Maximum stretch/hormone concentrations reported the optimized values that map to maximum network weights and define the linear transfer functions (Eq. 7-10, Fig. 3E-H).

| | **Determined from Network Model** | | **Calculated by compartmental model/from literature** | **Optimized** |
|---|---|---|---|---|
| | Baseline network weight | Maximum network weight | Baseline stretch/ hormone concentrations | Maximum stretch/ hormone concentrations |
| **Stretch (Eq. 7)** | | $w_{\text{stretch,max}} = 0.6$ | $\max(F_{e,f})_{\text{baseline}} = 2.23$ | $\max(F_{e,f})_{\text{max}} = 3.18$ |
| **AngII (Eq. 8)** | | $w_{\text{AngII,max}} = 0.6$ | $c_{\text{AngII,baseline}} = 4.40$ [pg/ml] (Ruzicka et al. 1995) | $c_{\text{AngII,max}} = 69.1$ [pg/ml] |
| **E2 (Eq. 9)** | $w_{\text{baseline}} = 0.1335$ | $w_{\text{E2,max}} = 1$ | $c_{\text{E2,baseline}} = 4.71$ [pg/ml] (Rosenblatt 1988; Jankowski et al. 2001) | $c_{\text{E2,max}} = 178$ [pg/ml] |
| **P4 (Eq. 10)** | | $w_{\text{P4,max}} = 0.85$ | $c_{\text{P4,baseline}} = 2.98$ [ng/ml] (Morishige et al. 1973; Blair and Mickelsen 2006) | $c_{\text{P4,max}} = 659$ [ng/ml] |

### 1. Sensitivity analysis for the multiscale model calibration

To test how well the optimized linear transfer function parameters ($\max(F_{e,f})_{\text{max}}$, $c_{\text{AngII,max}}$, $c_{\text{E2,max}}$, $c_{\text{P4,max}}$) constrained our growth predictions, we conducted a sensitivity analysis by increasing and decreasing the individual parameters from the optimized values. Supplemental Figure 2 demonstrates how changing the optimized parameters affects the error between model predictions and experimental growth data. Our sensitivity analysis indicates that the optimized parameters lead to minimum error for all simulated conditions. As expected, VO growth predictions are most sensitive to $\max(F_{e,f})_{\max}$, AngII infusion growth predictions are most sensitive to $c_{\text{AngII,max}}$, E2 infusion growth predictions to $c_{\text{E2,max}}$, and P4 infusion growth predictions to $c_{\text{P4,max}}$.

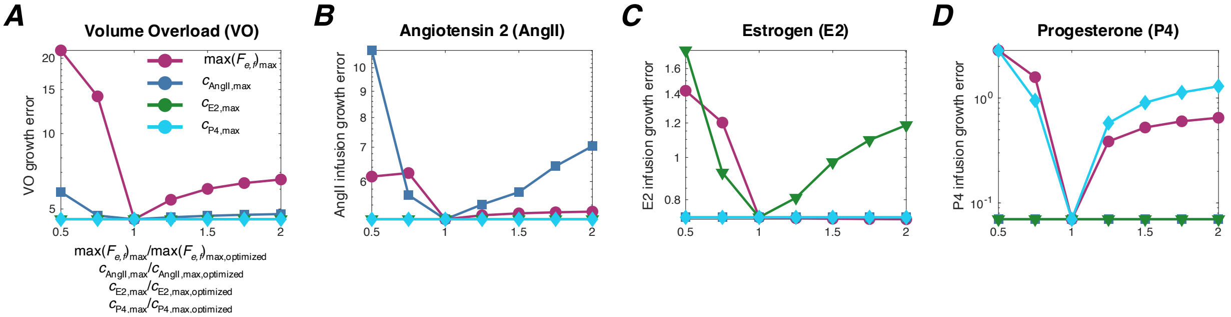


**Supplemental Figure 2: Coupling sensitivity analysis.** The optimized maximum stretch ($\max(F_{e,f})_{\max}$) and hormone concentrations ($c_{\text{AngII,max}}$, $c_{\text{E2,max}}$, $c_{\text{P4,max}}$) were varied 50% to 100% from the optimized values (Suppl. Table 2) and the resulting error in growth due to experimental A.) VO and infusion of B.) AngII, C.) E2, and D.) P4 were calculated. As the plots demonstrate, the optimized parameters lead to minimum error.
